# Interaction between facial expression and color in modulating ERP P3

**DOI:** 10.1101/2024.09.16.612994

**Authors:** Yuya Hasegawa, Hideki Tamura, Shigeki Nakauchi, Tetsuto Minami

## Abstract

The relationships between facial expression and color affect human cognition functions such as perception and memory. However, whether these relationships influence attention remains unclear. Additionally, whether facial expressions affect selective attention is unknown; for example, reddish angry faces increase negative social evaluation or emotion intensity, but it is unclear whether selective attention is similarly enhanced. To investigate these questions, we examined whether event-related potentials for faces vary depending on facial expression and color by recording electroencephalography (EEG) data. We conducted an oddball task using stimuli that combined facial expressions (angry, neutral) and facial colors (original, red, green). The participants counted the number of times a rarely appearing target face stimulus appeared among the standard face stimuli. The results indicated that the difference in P3 amplitudes for the target and standard faces depended on the combinations of facial expressions and facial colors; the P3 amplitudes for red angry faces were greater than those for red neutral faces. Additionally, there was no significant main effect or interaction effect of facial expression or facial color on P1 amplitudes for the target, and there were significant main effects of facial expression only on the N170 amplitude. These findings suggest that the intensity of a human’s selective attention to facial expressions varies according to the higher-order semantic processing of the relationship between emotion and color rather than simple facial expression or facial color effects individually. Our results support the idea that red color increases the human response to anger from an EEG perspective.

**Significance Statement:** It remains unclear whether selective attention to faces is modulated by the relationships between facial expression and color. Using an oddball task and recording EEGs, we showed that the event-related potentials reflecting selective attention are modulated by the interaction between facial expression and facial color, although the interaction was not found at earlier ERP stages. These findings suggest that the intensity of selective attention to facial expressions is influenced more by the interaction between facial expression and facial color than by facial expression or facial color alone and that the interaction occurs as a higher-order processing step than facial expression or color recognition. Our results provide EEG evidence supporting the idea that red color increases the human response to anger.

## Introduction

Facial color affects the judgment of facial expressions, with reddish faces easily judged as an angry face and perceived as having a greater emotional intensity of anger (Kato et al., 2022; Minami et al., 2018; Nakajima et al., 2017; Peromaa & Olkkonen, 2019; Thorstenson, Pazda, et al., 2021). Additionally, facial color has been shown to influence perceived social characteristics such as friendliness, aggression, and health (Thorstenson & Pazda, 2021). Moreover, the effects of the relationship between facial expression and facial color are similar not only for real faces but also for emoticons, facial models and implicit faces; this relationship is also known to change with background color, suggesting that specific colors enhance emotion perception (Liao et al., 2018; Minami et al., 2018; Nguyen et al., 2023; Qin, 2021; Thorstenson, McPhetres, et al., 2021). Recent studies have focused on the relationship between facial expression and facial color memory and suggested that facial expression biases facial color memory (Hasegawa et al., 2024; Thorstenson, Pazda, et al., 2021). Thus, the relationship between facial expression and facial color, especially the relationship between anger and red, affects cognitive function, such as human judgment and memory. However, it is still unclear whether this relationship affects attention, which is another cognitive function. In the relationship between facial expression and attention, angry and fearful faces are known to bias visual attention, and it is believed that perceived threats may be among the factors that capture human attention (Fox et al., 2002; Mathews et al., 2003; Öhman et al., 2001). On the other hand, regarding the effects of facial color, previous studies have confirmed that making angry faces redder than usual increased negative social evaluation (e.g., aggressiveness and dominance) and the perceived emotional intensity (Thorstenson, McPhetres, et al., 2021; Thorstenson & Pazda, 2021). Therefore, the following questions arise: does a reddish angry face increase perceived threat and emotional intensity, and is the observer’s attention more biased toward a reddish angry face than toward other faces?

One of the measures for comparing human attention is the event-related potential (ERP) P3. P3 is an ERP component that occurs approximately 300 to 500 milliseconds after a presented stimulus at the parietal lobe as the third positive deflection; P3 is also known to reflect selective attention. The oddball paradigm is a task that induces a large P3 amplitude. In the oddball task, participants count how often a low-frequency or specified stimulus appears among stimuli at other frequencies. Large P3 amplitudes are elicited by low-frequency stimuli during the oddball task. With respect to the relationship between facial expressions and P3 amplitudes, previous studies have shown that attention to angry or fearful faces increases the P3 amplitude (Chai et al., 2012; Kiss & Eimer, 2008; Lin et al., 2020; Rossignol et al., 2005). Thus, angry faces affect both the P3 amplitude and attention. However, it is still unclear whether the addition of red color, which is strongly associated with anger, increases P3 amplitudes and selective attention.

Therefore, this study aimed to clarify the effect of the relationship between facial expression and facial color on selective attention. The P3 amplitude, which reflects selective attention, varies in magnitude depending on the intensity of attention to the target. Therefore, we hypothesized that there is an interaction effect of facial expression and facial color on attention when strong attention is caused by the relationship between anger and red rather than a color effect alone. For example, the P3 amplitude for red angry faces is larger than that for red neutral faces, and this trend is more pronounced than when angry faces with normal facial color are compared with neutral faces with normal facial color. In this study, we recorded participants’ electroencephalography (EEG) during an oddball task to investigate the effects of facial expression and facial color on human attention. Then, we compared the P3 amplitudes to estimate how differences in facial expression and facial color influence attention.

## Method

### Participants

Twenty Japanese students (5 women and 15 men, mean age = 22.50 ± 1.00 years) at Toyohashi University of Technology participated in the experiment. The sample size was calculated using PANGEA (Westfall et al., 2014) with an effect size of *d* = 0.4, *power* = 0.8, and *α* = 0.05, and we found that 19 participants were needed. Assuming a possibility of data rejection due to any EEG artifacts, we recruited 20 participants. Before joining the experiment, the participants were provided with an introduction to the experiment, excluding the study’s hypothesis, and gave informed consent. All participants had normal color vision, as verified by the Ishihara Color Vision Test Chart II Concise Version 14 Plate (the Public Interest Incorporated Foundation Isshinkai, Handaya Co., Ltd., Tokyo, Japan). This experiment was conducted with the approval of the Ethics Committee for Human Research at Toyohashi University of Technology and adhered strictly to the approved guidelines of the committee and the Declaration of Helsinki. This study was not preregistered.

### Stimuli

Facial stimuli (Figure 1) were angry and neutral faces of two Japanese individuals (one woman and one man) obtained from the ATR Facial Expression Image Database (ATR-Promotions, Kyoto, Japan, https://www.atr-p.com/products/face-db.html). The hair, ears, and neck in the images were removed using Photoshop (Adobe Systems Inc., San Jose, CA, USA), with the edited image presenting an oval shape. All images were adjusted to maintain an average image luminance of 16.9 cd/m^2^ using SHINE_color, a MATLAB 2021a toolbox (Ben, 2021; Willenbockel et al., 2010). Based on an experiment by Nakajima et al. (2017), we created colored facial stimuli by manipulating the a* (red– green) value in CIE L*a*b* (Nakajima et al., 2017). There were three facial color conditions: original (no manipulation), red (a*+12 units), and green (a*-12 units). In total, twelve image stimuli (2 individuals × 2 facial expressions × 3 facial colors) were prepared. The image dimensions were 4.2° × 5.5°. The mean and standard deviation of the CIE L*a*b* values for the original faces were L*= 47.60 ± 0.06, a*= 7.61 ± 0.09, and b*= 22.41 ± 0.06. The background color was always gray ([*x, y*] = [0.31, 0.35], Y = 17.09 cd/m^2^).

**Figure 1.**
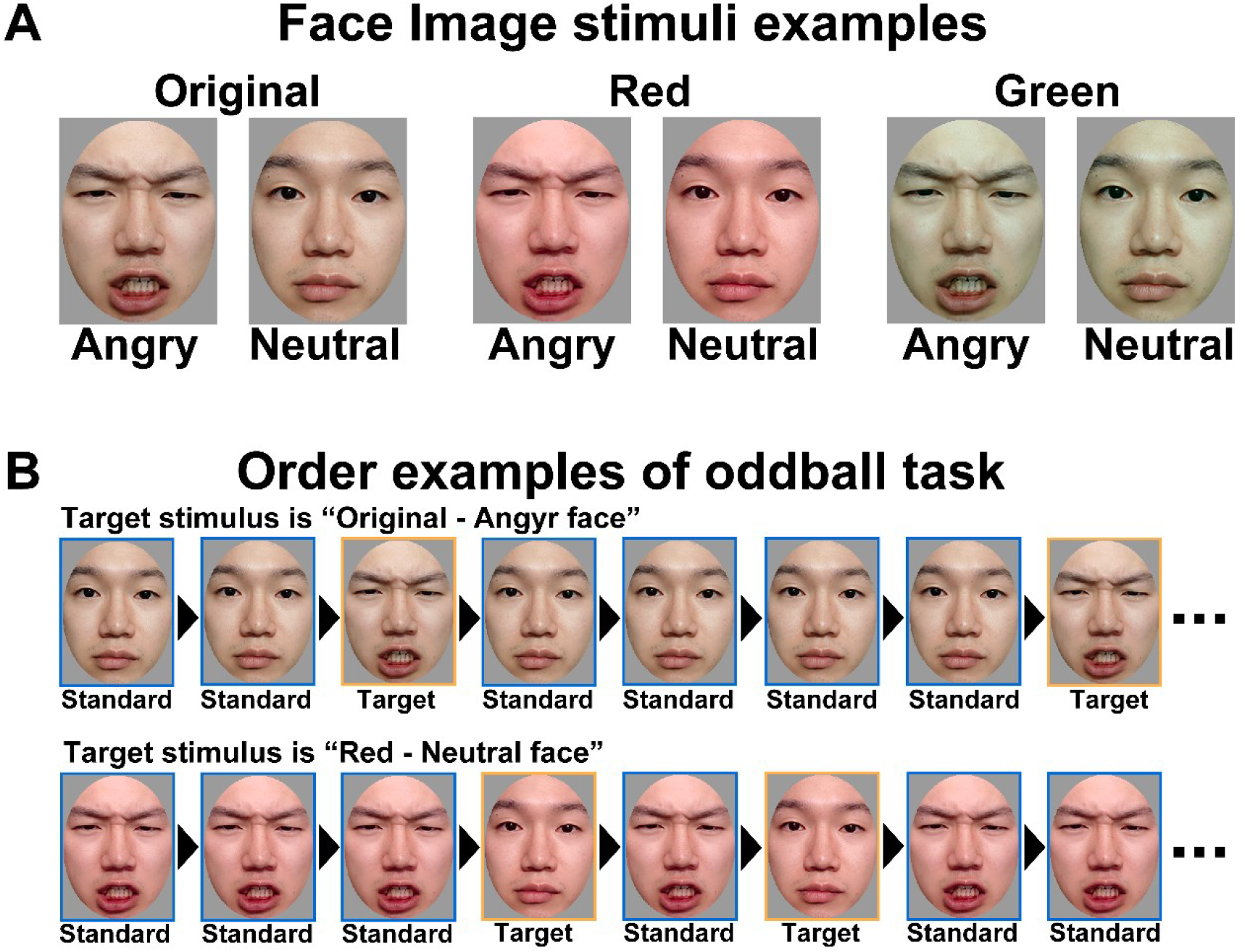
A: Examples of the image stimuli for each condition. There were two facial expression conditions, angry and neutral, and three facial color conditions, original (no manipulation), red (a*+12 units), and green (a*-12 units). Two face models (1 woman and 1 man) were used for each the six conditions (2 facial expression × 2 facial color). B: Examples of presentation order for oddball task. Standard stimuli are presented at high frequency, and target stimuli are presented at low frequency. The faces in the figure are from one of the authors (Y.H.) and were not used in the experiment.

### Apparatus

The experiment was conducted in a dark magnetically shielded room. The stimuli were presented on a monitor (VIEPixx/EEG, VPixx Technologies Inc., Canada; resolution: 1920 × 1080; frame rate 120 Hz). The white point of the monitor was [*x, y*] = [0.30, 0.33], *Y* = 91.23 cd/m^2^. The participants were seated and performed the task while keeping their heads on a chin rest positioned 60 cm away from the display. Psychotoolbox 3.0.17 served as the experimental control software (Brainard, 1997; Kleiner et al., 2007; Pelli, 1997). EEG data were acquired by 64 channels of electrodes and 6 channels of external sensors at a sampling frequency of 512 Hz using BioSemi ActiveTwo (Amsterdam, Netherlands) and recorded by the ActiveTwo System.

### Procedure

In the experiment, we used 6 image stimulus pairs. Three pairs were prepared for each face stimulus (2 persons): the original color angry face and original color neutral face, the red angry face and red neutral face, and the green angry face and green neutral face. An oddball task was performed with the high-frequency stimulus as the standard stimulus and the low-frequency stimulus as the target stimulus in these pairs. In each trial, when the target stimulus was a red angry face, the standard stimulus was a red neutral face, and when the target stimulus was a red neutral face, the standard stimulus was a red angry face. Therefore, a participant performed 12 trials (6 pairs × 2 targets) of the oddball task in total. Standard and target stimuli were presented in a random order during each trial. The frequency of standard and target stimuli was always Standard : Target = 1 : 4, and the target stimulus was presented 10∼15 times. The participants were asked to count the number of times the target stimulus appeared. Figure 2 shows a summary of the experimental procedure. First, a task description was presented until the participant pressed the enter key. After a 1.0 second interstimulus interval, the fixation points and facial expression stimuli (standard or target stimulus) were presented repeatedly at 0.5 second intervals. After the end of the presentation, the participant recorded the number of times the specified facial stimulus (target stimulus) appeared using a numeric keypad. The experiment was conducted in two blocks, one for counting angry faces and one for counting neutral faces (6 trials per block), and the order of the blocks differed among the participants. The order of the 6 pairs (2 persons × 3 facial color) presented within a block was random. The participants could take breaks between trials and blocks. The participants wore EEG equipment throughout the experiment, and their EEGs were recorded during the task. Additionally, to reduce motion noise, the participants were instructed to avoid counting with their fingers or voices.

**Figure 2.**
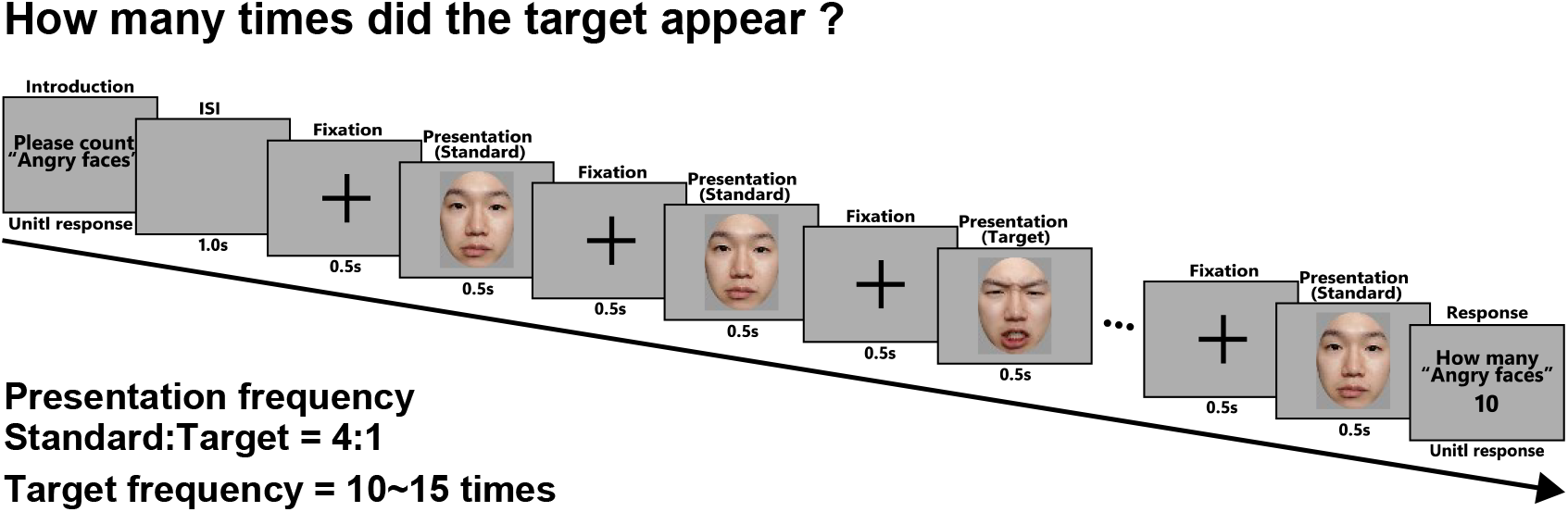
Procedure of experiment when the target stimuli are angry faces. In the repetition presentation phase, the stimuli were presented in a random order with a frequency of Target : Standard = 1 : 4. After the presentation phase, the participant recorded the number of times the specified facial stimulus (target stimulus) appeared in the instruction phase using a numeric keypad. The ratios of text screens, fixation cross, and stimuli depicted in this figure differs from the actual ratio.

### Preprocessing of EEG data

For preprocessing, the EEG data were downsampled to 200 Hz, and a high-pass filter (1 Hz) and the function “cleanLineNoise” in EEGLAB were applied to eliminate line noise such as white noise, power supply noise (60 Hz) and its harmonic frequencies (120, 180, and 240 Hz). The significance cutoff level was *p* = .01. Additionally, electrodes that did not measure the data well were removed using the function “clean_rawdata” in the EEGLAB tool (unchanged interval: 5 seconds, correlation with surrounding electrodes: less than 0.85, far from average: 4 times the standard deviation, removal using the ASR algorithm). The remaining data were subsequently interpolated by the spherical spline interpolation method. Moreover, artifact removal was performed by eliminating ocular components using adaptive mixture independent component analysis (AMICA) and ICLabel (Leutheuser et al., 2013; Pion-Tonachini et al., 2019). Finally, we set the time of stimulus presentation as 0 ms and extracted the EEG data at −100∼1000 ms. One participant was excluded from the analysis because all EEG data of target stimuli for one condition were excluded by this preprocessing.

### Statistical analysis

#### P3

First, we extracted the channel-average EEG at Cz, CPz, CP1, CP2, and Pz from the preprocessed EEG. The baseline EEG was the average EEG of −100∼0 ms . Next, the mean amplitudes at 300∼500 ms after the presentation of the standard and target stimuli were calculated for each trial. Then, the mean amplitude of the standard stimulus was subtracted from the mean amplitude of the target stimulus during the same trial, and we calculated the average for each of the facial expression (anger, neutral) and facial color (original, red, green) conditions (for a total of 6 conditions). Quartile range exclusion was subsequently performed for each condition to exclude outliers, and the data of one participant were excluded. Therefore, the final statistical analysis included 18 participants.

We first conducted the Shapiro-Wilk test to confirm that the data were normally distributed. If the data were confirmed to be normal, we performed a repeated measure two-way analysis of variance using R. If the data did not follow a normal distribution, we performed a nonparametric test with nparLD, an R software package (Noguchi et al., 2012). ANOVA-type statistics (ATS) were calculated using nonparametric tests. Moreover, the p values underwent post hoc correction by the Holm method.

#### N170 (Left, Right)

We extracted the channel-average EEG at 5 channels near the temporal area (Left: TP7, P5, P7, P9, PO7; Right: TP8, P6, P8, P10, PO8) from the preprocessed EEG. The baseline time window was the same as that for the P3 amplitude. Then, the peak amplitudes at 150∼200 ms after the presentation of the target stimuli were calculated for each trial, and we averaged them for each condition using the same method as that used for P3. Afterward, the same statistical analysis as that used for P3 was performed on the left and right peak amplitudes. We used the same participant data as those used in the statistical analysis of P3.

#### P1

We extracted the channel-average EEG at Iz, Oz, O1, O2, and POz from the preprocessed EEG. The baseline time window was the same as that for the P3 and N170 amplitudes. The P1 amplitude was calculated using the same procedure used for N170, except that the time window of the peak amplitude was 80∼120 ms. Afterward, the same statistical analysis as that used for P3 was performed on the left and right peak amplitudes. We used the same participant data as those used in the statistical analysis of P3.

## Results

### P3

Figure 3 shows the average wave for the target and standard stimuli for each condition. Figure 4 shows the mean P3 amplitude for each facial expression and facial color condition. We found a significant main effect of facial expression (*F*(1,17) = 10.089, *p* < .01, *η*_*p*_^2^ = 0.372) and a significant interaction effect between facial expression and facial color (*F*(1.97,33.48) = 3.747, *p* < .05, *η*_*p*_^2^ = 0.181). The post hoc results revealed that the P3 amplitude for angry faces was greater than that for neutral faces (*t*(17) = 3.176, *p* < .01, Cohen^′^s *d* = 0.749), and the P3 amplitude for red angry faces was greater than that for red neutral faces (*t*(17) = 3.382, *p* < .05, Cohen^′^s *d* = 0.797).

**Figure 3.**
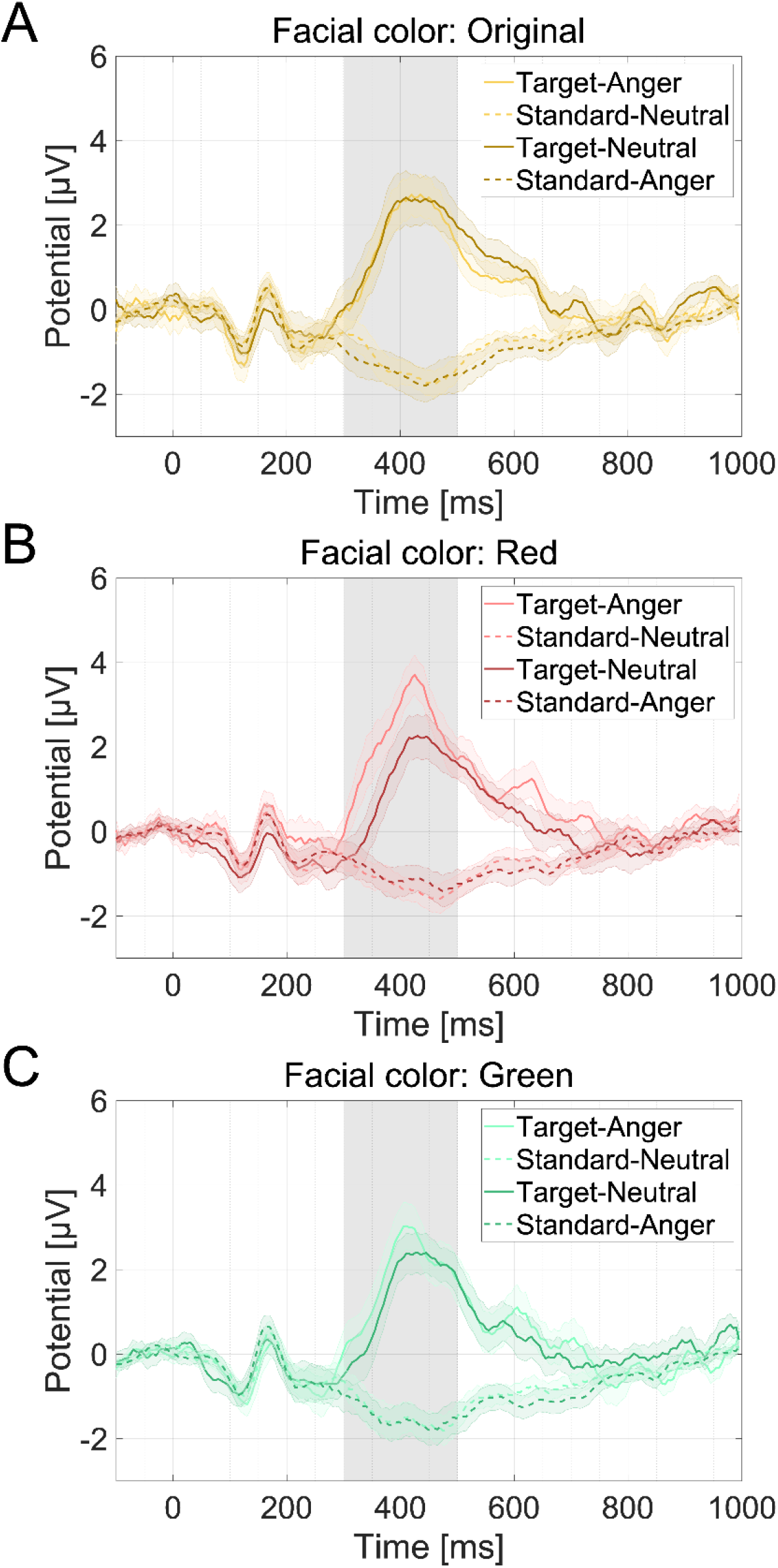
Channel-average EEG waves of mean for each condition (A: Original, B: Red, C: Green). The bands covering the line and the dashed line represent the standard error of the mean. The gray bands at 300∼500 ms are the time windows of P3 analysis. The data were smoothed for plotting and were not used in the analysis.

**Figure 4.**
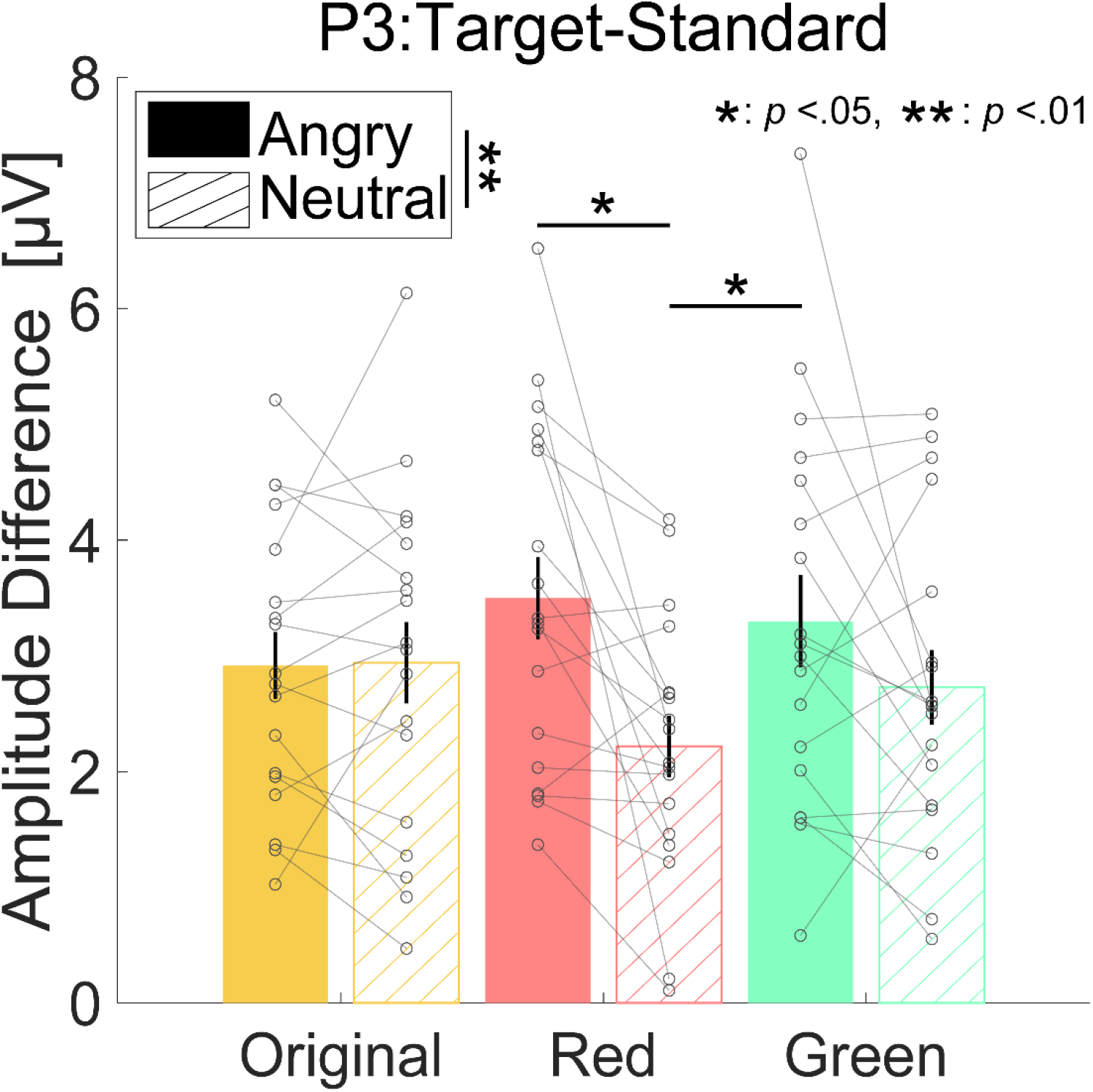
Mean of the difference in P3 amplitude [μV] for target and standard stimuli. Each gray points represent individual data. The color of each bar and the label on the horizontal axis indicate facial conditions. The filled bars indicate that target stimuli were angry faces, and the hatched bars indicated that target stimuli were neutral faces. Error bars show the standard error of the mean.

### N170

Figure 5 shows the mean N170 amplitude for each facial expression and facial color condition. We found a significant main effect of facial expression on both the left and right sides (Left: *ATS*(1) = 6.940, *p* < .01; Right: *F*(1,17) = 10.100, *p* < .01, *η*_*p*_^2^ = 0.373). Post hoc tests revealed that the N170 amplitudes for angry faces were greater than those for neutral faces (Left: *Z*(17) = −2.896, *p* < .01, *r* = 0.683 ; Right: *t*(17) = −3.178, *p* < .01, Cohen^′^s *d* = 0.749). These results are similar to those of previous studies, which revealed larger N170 amplitudes for negative facial expressions than for neutral facial expressions.

**Figure 5.**
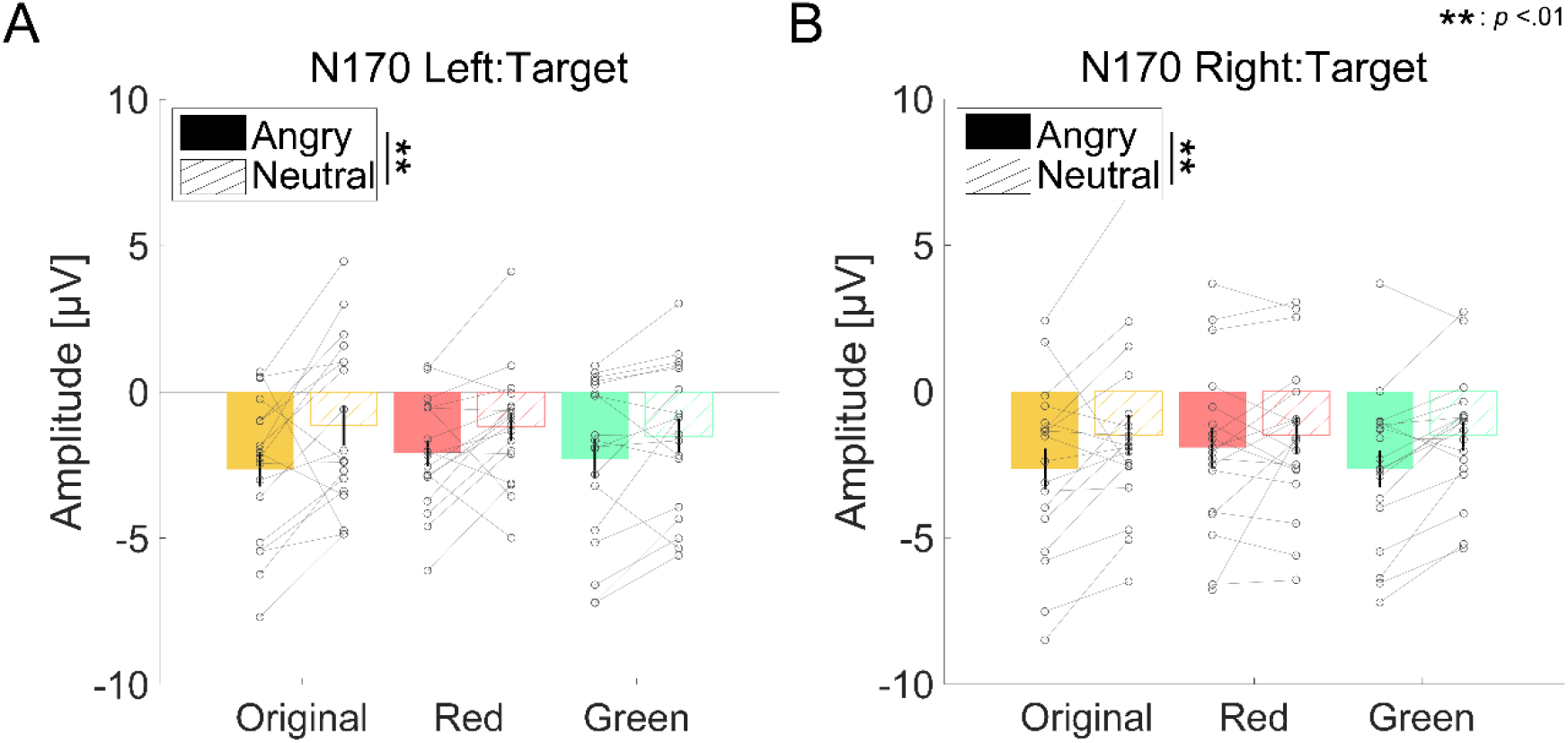
Mean of the N170 amplitude [μV] for target stimuli (A, Left side; B, Right side). The color of each bar and the label on the horizontal axis indicate facial conditions. The filled bars indicate that target stimuli were angry faces, and the hatched bars indicate that target stimuli were neutral faces. Error bars show the standard error of the mean.

### P1

Figure 6 shows the mean P1 amplitude for each facial expression and facial color condition. There was no significant main effect or interaction effect of facial expression and facial color (expression: *F*(1,17) = 0.065, *p* = .802, *η*_*p*_^2^ = 0.004 ; color: *F*(1.85, 31.53) = 1.051, *p* = .357, *η*_*p*_^2^ = 0.058 ; interaction: *F*(1.52, 25.90) = 3.107, *p* = .074, *η*_*p*_^2^ = 0.155).

**Figure 6.**
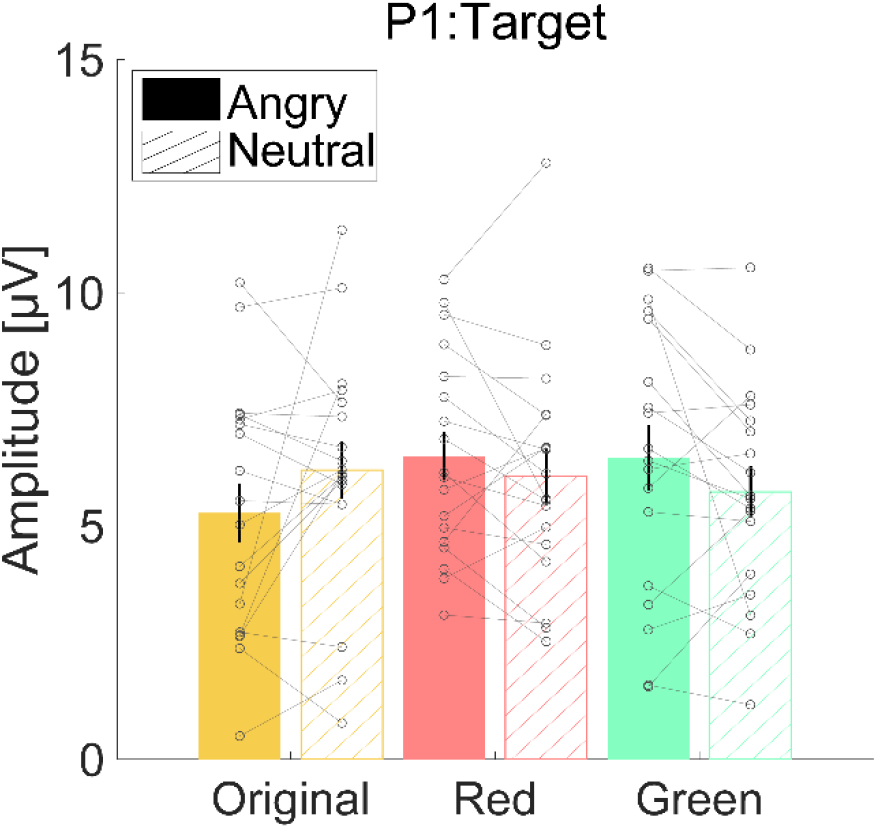
Mean P1 amplitude [μV] for target stimuli. The color of each bar and the label on the horizontal axis indicate facial conditions. The filled bars indicate that target stimuli were angry faces, and the hatched bars indicate that target stimuli were neutral faces. Error bars show the standard error of the mean.

## Discussion

In this study, we used an oddball task to investigate whether the relationship between facial expression and facial color influences selective attention and recorded participants’ EEGs during the task. The results revealed that the P3 amplitude for red angry faces was greater than that for red neutral faces, suggesting that the selective attention given to red angry faces was greater than that given to red neutral faces. Moreover, there was no main effect or interaction effect of facial expression and facial color for the P1 amplitude and only a main effect of facial expression for the N170 amplitude. On the other hand, there was a main effect of facial expression and an interaction effect between facial expression and facial color for the P3 amplitude. These results indicate that the relationship between facial expression and facial color is represented by higher-order processing along the P1, N170, and P3 time axes. P1 amplitudes reflect the initial attentional processing of stimuli, and N170 amplitudes reflect differences in facial expression (Hillyard & Anllo-Vento, 1998; Hinojosa et al., 2015). In contrast, P3 amplitudes reflect higher-order cognitive processing, such as conscious attention (Polich, 2007). Therefore, the results of this study suggest that the intensity of a human’s selective attention to facial expressions varies according to the higher-order semantic processing of the relationship between emotion and color rather than facial expression or facial color effects alone.

Facial stimuli that evoke emotions, such as anger, selectively bias attention (Öhman et al., 2001). Previous studies have reported that intensifying the redness of angry faces increased perceived emotional intensity, aggression, and threat (Thorstenson, McPhetres, et al., 2021; Thorstenson & Pazda, 2021). The results of this study support the idea that red increases the human response to anger from an EEG perspective.

In addition, the P3 amplitude, which reflects selective attention, is associated with memory. P3 is also an indicator of the degree of encoding and recall, and previous studies have suggested that a high P3 amplitude indicates the importance of encoding and the degree of successful recall (Fabiani et al., 1986; Karis et al., 1984; Polich, 2007). Emotionally relevant stimuli are known to be more strongly anchored in memory than are neutral stimuli, and the results of this study suggest that red angry faces have a greater influence on human memory (Dolcos & Cabeza, 2002). These results support previous studies that suggest that facial color memory for angry faces is biased toward more reddish and yellowish colors than that for actual facial color or neutral faces (Hasegawa et al., 2024; Thorstenson, Pazda, et al., 2021).

Contrary to expectations, angry faces with a green facial color, which is the color opposite to red, also presented high P3 amplitudes. The contextual incongruence of the green color with the angry face may have caused a high cognitive load, which affected the P3 amplitude. Minami et al. (2009) reported increased P3 amplitudes in the oddball task for unnatural stimuli such as inverted faces and deviant color faces and suggested that asymmetry in P3 amplitudes reflects the unnaturalness of visual stimuli (Minami et al., 2009). Therefore, the reason why the P3 amplitude for green angry faces was not lower than that of red angry faces, contrary to expectations, is believed to be due to the unnaturalness of an angry face having a green facial color.

The results of this study revealed that the N170 amplitude differed depending on facial expression, with the N170 amplitudes for angry faces being larger than those for neutral faces. These results support the findings from previous studies that emotional relatedness increases the N170 amplitude during facial processing (Hinojosa et al., 2015). On the other hand, there were no differences in the N170 amplitude across different facial colors. It has been reported that the N170 amplitude reflects facial color processing and that the amplitude increases with respect to the unnaturalness of facial color (Minami et al., 2011; Nakajima et al., 2012). The stimuli used in this experiment had a color change of a*±12 units, which is considered not unnatural as a facial color. Thus, our results suggest that the a*±12 level of facial color change does not affect the N170 amplitude. This finding is also consistent with the results of Nakajima et al. (2012), where the N170 amplitude when the hue angle was changed in the red direction (−45° when the original color was 0°) did not differ from the N170 amplitude for the normal facial color (Nakajima et al., 2012).

The limitations of this study are as follows. First, the individual characteristics of the participants were not researched in this experiment. The ability to detect faces, recognize or process facial expressions, and bias attention toward facial expressions such as angry faces varies depending on trait anxiety, autism spectrum disorder or Moebius syndrome (Golarai et al., 2006; Quettier et al., 2023; Surcinelli et al., 2006; Tanaka & Sung, 2016; Telzer et al., 2008). Therefore, it is possible that the individual characteristics of participants may have affected differences in attention to facial expressions, and it is necessary to examine the differences in the magnitude of effects due to individual characteristics in the future.

Second, all participants in this experiment were Japanese, and the facial stimuli used were also Japanese models. Color preferences for emotion and facial color are known to vary across cultures, and the association between emotion and color is known to be developmentally variable (Boyatzis & Varghese, 1994; Han et al., 2018; Jonauskaite et al., 2020). Hence, it is appropriate to interpret the findings of this study as being based on phenomena observed under specific conditions, and their validity is limited within certain populations.

## Conclusion

We investigated whether ERP P3 varies with facial expression and facial color according to the hypothesis that humans bias more selective attention to red angry faces than to neutral facial expressions or original facial colors. The results revealed an interaction effect between facial expressions and facial colors on P3, which was not observed in early responses such as P1 and N170, and the P3 amplitudes for red angry faces were greater than those for red neutral faces. These findings indicate that humas bias more attention to red angry faces than to red neutral faces and suggest that the intensity of attention to human facial expressions does not depend on facial expressions or facial color effects alone but on the higher-order semantic relationship between emotion and color. Our findings support the idea that red increases or biases the response to anger from an EEG perspective.

## Acknowledgment

This work was supported by JSPS KAKENHI (Grant Numbers JP22K1789 to H.T., JP20H05956 to S.N., JP20H04273 to T.M., and JP23KK0183 to T.M.), JST SPRING, Japan (Grant Numbers JPMJSP2171 to Y.H.), and the student fellowship program for the Leading Graduate School at Toyohashi University of Technology to Y.H.

Commercial relationships: None.

## Author contributions

T.M. and Y.H. designed the research; T.M., Y.H. and H.T. provided experimental code and conducted the analysis; T.M., Y.H. and H.T. wrote the paper; S.N. provided feedback on the manuscript. All the authors were responsible for funding acquisition.

## Data availability

The datasets generated and/or analyzed during the current study are available from the corresponding author upon reasonable request.

